# Sequestration of oxidative is necessary but not sufficient enough to conclude dopaminergic neuroprotective efficacy of curcumin: Insights from ALSS *Drosophila* Parkinson’s disease model

**DOI:** 10.1101/2024.09.19.613867

**Authors:** Limamanen Phom, Priyanka Modi, Abhik Das, Sarat C. Yenisetti

## Abstract

Turmeric is a centuries-old ethnomedicine in Asia. Previously our laboratory demonstrated in the adult life stage-specific (ALSS) *Drosophila* model of Parkinson’s disease (PD) that Curcumin (K)-mediated dopaminergic (DAergic) neuroprotection is absent in the transition stage of adult life during which late-onset neurodegenerative disorders like PD sets-in, suggesting its limitation as a therapeutic agent. The present study demonstrates that K can sequester the enhanced levels of brain oxidative stress (OS) during both adult life phases i.e. health and transition stages but confers neuroprotection only during the health phase. However, literature reviews illustrate that efficacy of supposed therapeutic agents was asserted by their ability to sequester OS in only young PD animal models. In this context, it is important to point out that despite encouraging results in animal models, therapeutic efforts to target the general state of OS failed to retard PD progression. To understand this paradigm, we further investigated ALSS regulation of molecular players in the brain of the ALSS fly PD model and discovered that K-mediated differential modulation of adaptive stress response through *dFOXO* contributes to health phase-specific neuroprotection. These observations suggest that apart from the study of OS markers; it is essential to understand the ALSS regulation of molecular players. The synergistic influence of OS and the ALSS dysfunctional molecular networks could be responsible for the DAergic neurodegeneration in PD. The insights suggest that sequestration of OS by a therapeutic agent is necessary, but inadequate to conclude its neuroprotective efficacy and push it to the next phase of preclinical/clinical evaluation.

## 1. Introduction

Parkinson’s disease is the second most common neurodegenerative diseases, and it is characterized by degeneration of DAergic neurons in the *substantia nigra* (SN) of midbrain (1–3). The neuronal loss in this region leads to motor symptoms like bradykinesia, resting tremor, postural instability, gait impairment and non-motor symptoms (NMS) like impaired olfaction, constipation, depression, increased daytime sleep and rapid eye movement sleep disorder (4). By the time clinical symptoms such as resting tremors show up, about 70% of DAergic neurons are already degenerated, being the critical challenge in addressing pathophysiology of PD.

Oxidative Stress is classically defined as a state of redox imbalance caused by an excess formation of oxidants or a defect in antioxidants (5–8). Accumulation of free radicals and subsequent neurodegeneration in specific brain regions have been proposed as the underlying factors in neurodegenerative diseases such as Alzheimer’s and PD (9,10). Clinical and preclinical studies point out that neurodegenerative diseases like PD are characterized by increased level of OS biomarkers and decreased level of antioxidant defense biomarkers in the brain and peripheral tissues (9,11,12), emphasizing the role of oxidative stress in DAergic neurodegeneration.

Many laboratories have shown the DAergic neuroprotective effectiveness of natural compounds in PD models. Curcumin (K), 1, 7-bis (4-hydroxy-3-methoxyphenyl)-1, 6-heptadiene-3, 5-dione, is the main, active, yellow-colored natural polyphenolic compound. It is extracted from rhizomes of *Curcuma longa* L (turmeric), a plant in the ginger family (Zingiberaceae). K is widely used all over the world and extensively used in India and other Asian countries as a spice, food preservative, dye and ingredient of many traditional medicines (13,14). Various researches have demonstrated both in cell culture and animal models (15), K as a potent anti-oxidant (16–21), anti-inflammatory (22–24), anti-cancer agent (25–27), iron chelating agent (28–30) and neuroprotective compound (31–33). In addition, more than 440 clinical investigations have been performed using K (34). Thus K is one such potential candidate that can be explored for therapeutic approach to several human diseases including neurodegenerative disorder like PD.

There are about 100 research articles on *Drosophila* model of PD, implicating role of OS in neurodegeneration (source: Pubmed). Further, about 9000 articles on causative role of oxidative stress in neurodegenerative diseases are published suggesting a strong link between OS and neurodegeneration (35–44).

Several laboratories have shown the effective role of various therapeutic compounds that modulate the altered enzymatic functions arising out of OS in model organisms. They analyzed the neuroprotective properties of nutraceuticals using OS markers as a gold standard and suggest the neuroprotective efficacy of those compounds basing on their ability to sequester OS alone and recommended them for therapeutic approaches in NDD such as PD (45–52).

In the adult life stage-specific (ALSS) *Drosophila* model of sporadic PD, we have previously demonstrated that paraquat (PQ) treatment induced motor defects and depleted brain dopamine (DA) levels in both health and transition phases of adult life. However K confers DAergic neuroprotection only during health span of adult life and not during the transition phase, the adult life stage during which late onset neurodegenerative diseases such as PD sets in. This observation underlines the limitation of K as a neuroprotective agent in late onset NDD-PD (53,54).

In the present study, in order to understand the mechanistic insights into the adult life phase specific therapeutic propensity of K, its oxidative stress sequestration efficacy is assayed by quantifying the stress markers, antioxidant and neurotransmitter enzyme activities during health and transition phase of the adult life span in *Drosophila* model of PD. We found that K can mitigate the brain specific (all the assays are performed in fly brain tissue lysate) OS both during health and transition stage of *Drosophila,* suggesting the ability of K to sequester the OS. However, for first time previously we demonstrated that K confers DA neuroprotection only during health span, suggesting the critical limitation of OS sequestration based studies in determining the neuroprotective efficacy of nutraceuticals/therapeutic molecules (53,54). Importantly multiple studies relating to screening DA neuroprotective compounds were performed in the adult animal brain belonging to health span (PD is a late onset disease!). Hence it is important to underline the importance that sequestration of boosted OS alone is not enough to prevent the DAergic neurodegeneration in PD.

Hence, all these studies suggest that apart from assessment of OS, understanding the genetic/molecular mechanisms that could be synergistically responsible for the DAergic neurodegeneration in PD is essential to screen and develop novel therapeutic molecules for late onset NDDs such as PD. In *Drosophila*, adaptive stress response controlled by the *JNK-FOXO* axis has been greatly implicated in longevity, oxidative stress resistance (55–58). Further, K analogue mediated life span extension cum stress resistance in fly model has been demonstrated to require presence of *dFOXO* signaling (59). Therefore, in the present study K-mediated adult life stage specific (ALSS) differential modulation of *dFOXO* was also analyzed.

All these comprehensive studies illustrate that sequestration of OS may be essential, but alone not sufficient enough to promote DAergic neuroprotection, K-mediated adult life phase specific modulation of *dFOXO* and subsequent neuroprotection only in health phase and not during transition phase, together all these studies illustrates that sequestration of OS may be a necessity but not sufficient enough to confer DAergic neuroprotective efficacy and it is critical to look into the involvement of life-stage specific regulation of brain-specific molecular pathways and networks. This understanding is essential to explain the health phase specific dopaminergic neuroprotective efficacy of K and also develop successful DAergic therapeutic strategies for late-onset NDD such as PD.

## 2. Results

### 2.1. Curcumin diminishes PQ induced ROS level during the Health and Transition phase of adult life stage in fly model of PD

The modulatory efficacy of K on PQ-induced elevation in brain ROS levels was determined. It was found that in 4-day-old flies 34% enhanced level of ROS by PQ exposure was effectively diminished upon R_500µM_ and R_1mM_ K co-treatment (20% and 60% respectively). Also, in 55 day old flies there was 5 fold increase in ROS level by PQ treatment when compared to control, suggesting that at later stage of life, the organism becomes more susceptible to OS conditions. K co-treatment (R_500µM_ and R_1mM_) could reduce the elevated ROS levels by 60% and 35% respectively. PQ elevated the ROS levels in both the age group of flies and K *per se* did not have any detrimental effects on 4-day flies. However, in the 55 day old flies it was found that concentration of K *per se* (K_500µM_ and K_1mM_) increased the ROS level when compared to control (Fig.1A,B) suggesting that K itself may act as potential pro-oxidant and therefore the necessity to carefully screen the compound before prescribing them as a therapeutic agent.

**Figure 1.**
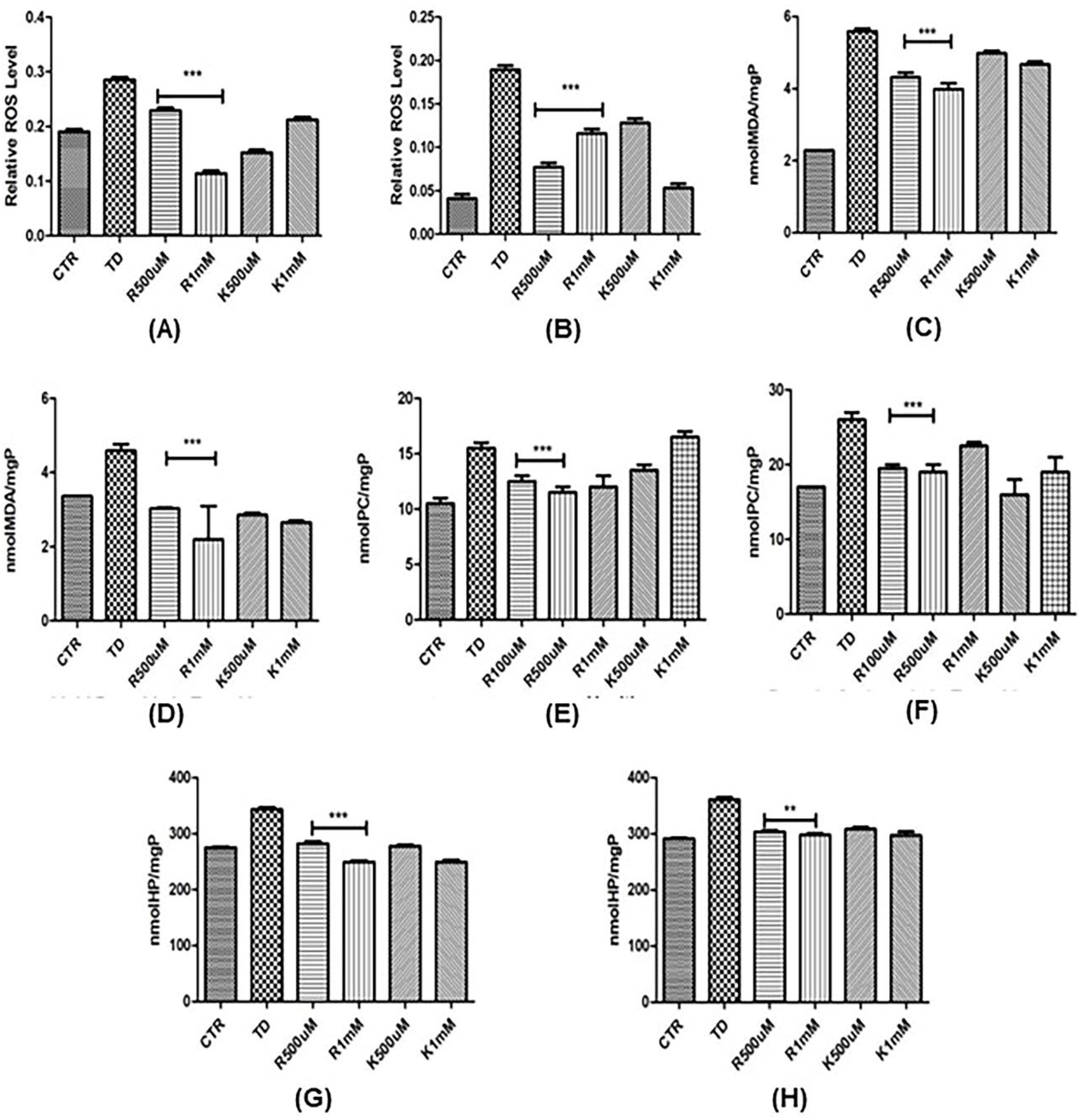
Estimation of antioxidant enzyme activities. **(A-H)**. Measurement of ROS in health (A) and transition (B) phase; Lipid peroxides in health (C) and transition (D); Protein carbonyls in health (E) and transition (F); Hydroperoxides in health (G) and transition (H); Irrespective of the age of flies, K co-treatment shows sequestration of oxidative stress in all the markers studied. (CTR=Control, TD=Treated with PQ, R(Rescue)= PQ+K, K=Curcumin *per se*)

### 2.2. Curcumin diminishes PQ induced Lipid peroxidation levels during Health and Transition phase of adult life stage in fly model of PD

To study the neuroprotective activity of K by estimating Lipid Peroxidation (LP), the levels of MDA which is an end product of LP was analyzed as one of the oxidative stress markers. PQ exposure upregulated LP by 60% and partially rescue was observed upon K treatment (about 23% and 31% reduction in R_500µM_ and R_1mM_ respectively) in health stage. In the transition stage, PQ treatment upregulated LP by 24% when compared to control. The increase in LP level was also partially rescued (about 30%) with both the K concentration used (R_500µM_ and R_1mM_) which indicate the effective free radical sequestration properties of K (Fig.1C,D).

### 2.3. Curcumin diminishes PQ induced Protein carbonyl level during Health and Transition phase of adult life stage in fly model of PD

K neuroprotective efficacy was also studied by estimating the PC level. While there was marked increase in PC upon PQ exposure (44% during health stage and 27% increase in transition phase), K co-treatment could effectively attenuate the PC level by all the K concentrations used in health stage. There was 18%, 28% and 20% reduction in PC levels with all the tested concentrations of K (R_100µM_, R_500µM_ and R_1mM_ respectively). At transition stage there was reduction by 25% (R_100µM_,), 24% (R_500µM_) and 20% (R_1mM_). (Fig.1E,F).

### 2.4. Curcumin diminishes PQ induced Hydroperoxide level during Health and Transition phase of adult life stage in fly model of PD

Hydroperoxidation was increased by 30% in health stage and 13% in transition stage on PQ exposure. K co-treatment decreased the HP level by 20% and 30% in health stage and 15% and 18% in transition stage with both R_500µM_ and R_1mM_ concentrations respectively (Figure.1G,H). This result further confirms the potent free radical scavenging activity of K. It is evident that K can actively sequester the OS level in both the health and transition stage of adult life stages of *Drosophila*.

### 2.5. Curcumin diminishes PQ induced SOD level during Health and Transition phase of adult life stage in fly model of PD

SOD, the first line of antioxidant defense system was elevated by 20% in health stage and 27% in transition stage (Fig.2A,B) by PQ treatment. K co-exposure significantly reduced the anti-oxidant enzyme activity. It was found that R_1mM_ concentration could bring down the elevated SOD activity by 40% in health stage and 20% in transition stage. It suggests lowered level of OS in the fly brain with K co-treatment. In transition stage, it was found that K *per se* was also found to actively sequester OS as indicated by decreased SOD activity when compared to control. We also quantified the brain expression levels of SOD1 and SOD2 during health and transition stages that shows both SOD 1and 2 protein levels reduced under PD condition, but K interference confers no rescue of SOD 1, whereas health phase specific rescue observed for SOD2 (Fig S1-B, C).

**Figure 2.**
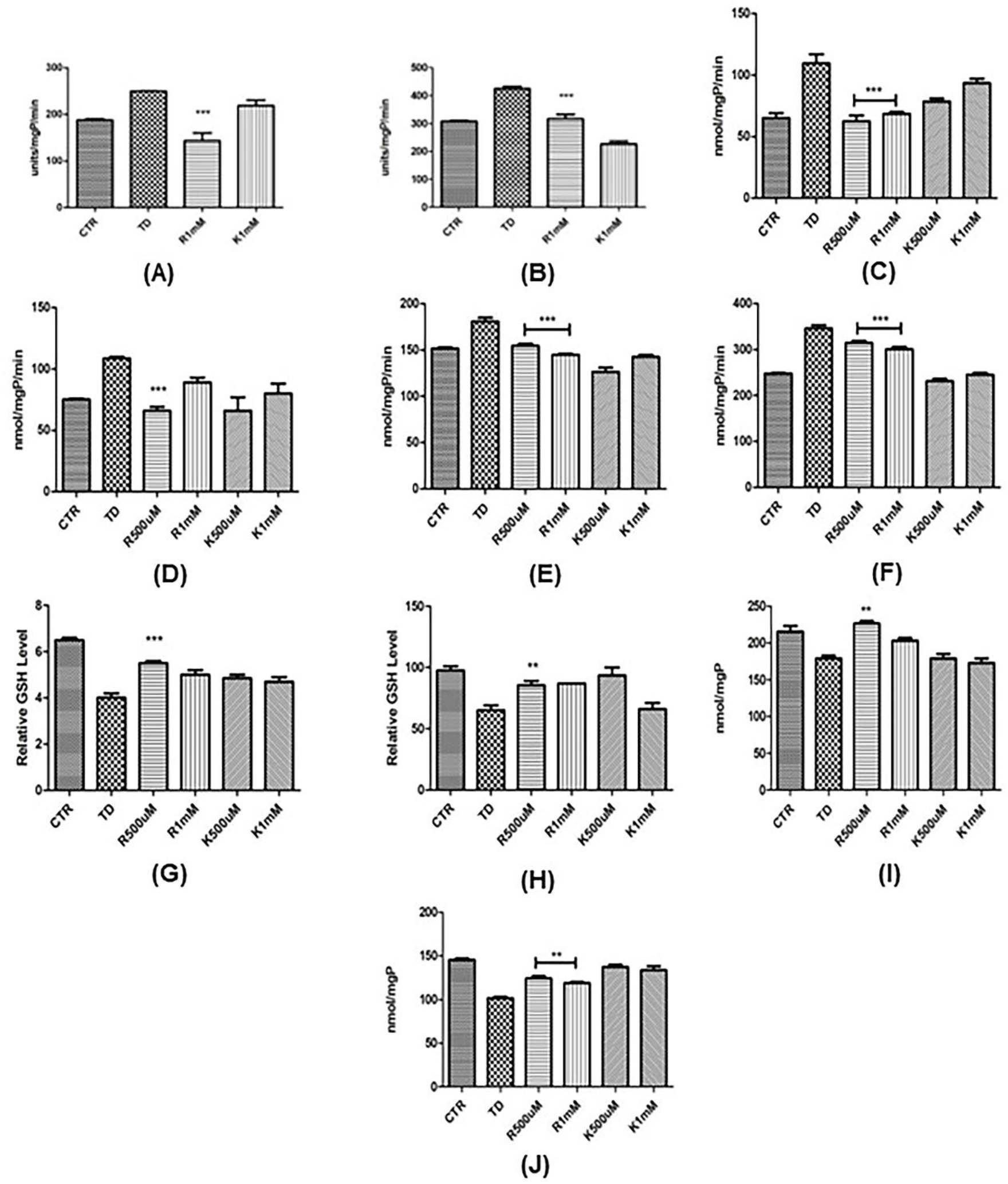
Quantification of non enzymatic stress markers. **(A-J)**. Estimation of SOD activities in health (A) and transition (B) phase; Catalase in health (C) and transition (D); GST in health (E) and transition (F); GSH level in health (G) and transition (H); Thiols in health (I) and transition (J); In all the enzyme activities measured, K co-treatment shows attenuation of the altered enzyme activities and levels triggered by PQ. (CTR=Control, TD=Treated with PQ, R (Rescue)= PQ+K, K=Curcumin *per se*)

### 2.6. Curcumin diminishes PQ induced CAT level during Health and Transition phase of adult life stage in fly model of PD

K effectively decreased the CAT activity which was elevated by PQ. Exposure of PQ increased the CAT level by 43% in health stage and 30% in transition stage (fig.2C,D). Co-treatment of K decreased the level by 44% and 35% in health stage and 40% and 15% in transition stage by both R_500µM_ and R_1mM_ respectively. While K *per se* was also found to increase the CAT activity by 30% in health stage, there was no such significance in transition stage. The result shows antioxidant activity of K which could reduce the elevated antioxidant enzyme activity suggesting lowered level of OS by K action. However, expression levels of brain catalase during health and transition stages shows a reduction of protein level (20 and 10% respectively) under induced PD condition and K confers partial rescue during health phase only (Fig S1-A).

### 2.7. Curcumin diminishes PQ induced GST level during Health and Transition phase of adult life stage in fly model of PD

The antioxidant property of K was evident even by the measurement of GST activity. It was found that PQ elevated GST level by 20% during health stage and 30% in transition phase when compared to controls (Fig.2E,F). The elevated level was decreased by 18% and 22% in health stage and 15% and 19% in transition stage in both the concentrations used (R_500µM_ and R_1mM_) respectively. This result further reaffirms the antioxidant property of K and its modulatory effect against PQ induced OS in fly model of PD.

### 2.8. Curcumin replenishes PQ induced diminished Glutathione level during Health and Transition phase of adult life stage in fly model of PD

PQ exposure led to significant reduction in GSH activities in both the age groups of flies. There was reduction of 30% during health stage and 27% during transition stage (Fig.2G,H) suggesting enhanced levels of OS in brain. K significantly elevated the diminished level of GSH in both the age groups. There was an upregulation of 20% and 15% in health stage and 23% and 27% in transition stage with R_500µM_ and R_1mM_ concentration respectively. These results further suggest the efficacy of K as an antioxidant therapeutic agent.

### 2.9. Curcumin replenishes PQ induced Total Thiol level during Health and Transition phase of adult life stage in fly model of PD

K could also partially rescue the depleted thiols levels caused by PQ treatment. In health stage 18% depletion due to PQ exposure and was rescued by K cotreatment with R_500µM_ and R_1mM_ concentration (14% and 12% respectively) whereas in transition stage 27% inhibition in thiols levels was rescued to 17% and 14% upon cotreatment with R_500µM_ and R_1mM_ concentration respectively (Fig.2I,J). It further supports the previous findings in other markers that K has anti-oxidative property and has the potential as a therapeutic agent in PD.

### 2.10. Curcumin replenishes PQ inhibited AChE activity during Health and Transition phase of adult life stage in fly model of PD

K also partially rescued the PQ induced marginal inhibition of AChE activity in both the age groups. In health stage 20% inhibition in AChE activity was rescued by 13% and 16% and in transition stage 16% inhibition was rescued by 15% and 8% in both the R_500µM_ and R_1mM_ concentrations respectively (Fig.3A,B). This result further suggests the neuromodulatory properties of K and its potential efficacy as a therapeutic agent in PD.

**Figure 3.**
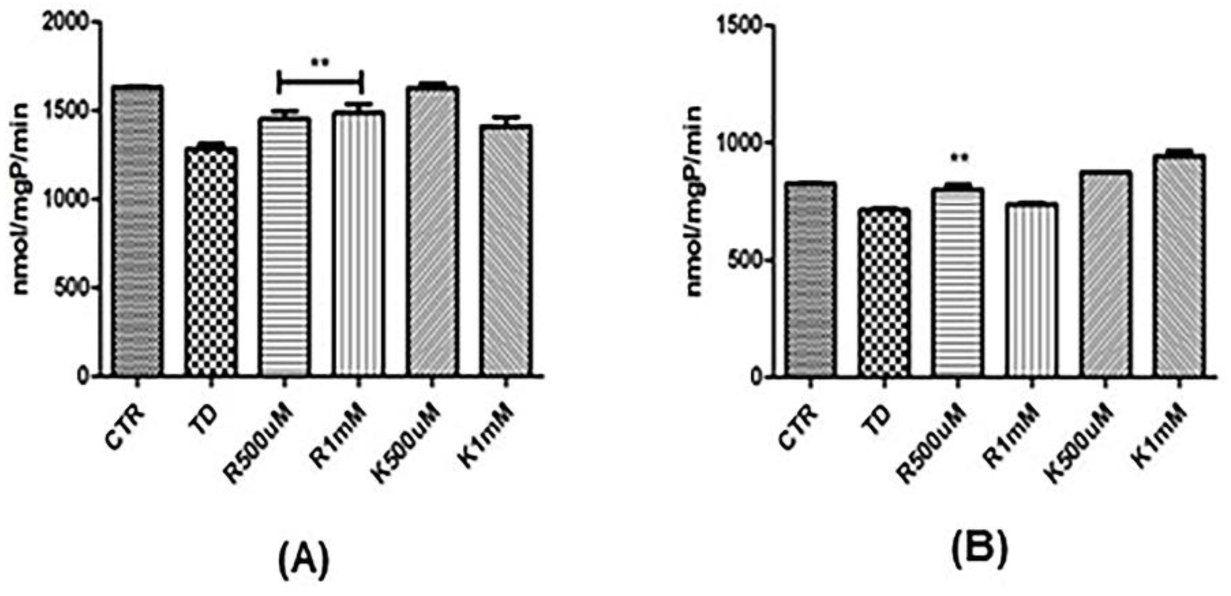
Estimation of Acteylcholine esterase activivity as a neurotoxicity marker. **(A,B)**. Measurement of AChE activity in health (A) and transition (B) stage. PQ inhibited AChE activity was rescued by K co-treatment in both the adult phase of Drosophila, suggesting the efficacy of K as a modulant against the impairment of neurotransmitter enzyme activity induced by PQ in fly model of PD; (CTR=control, TD=treated with PQ, Rescue (R)=PQ+K; K= curcumin *per se*)

### 2.11. *dFOXO* transcript expression level was diminished in the PD brain and K intervention rescues only during health phase

The downstream effect of *Bsk* signalling is brought about activation of transcription factor *dFOXO*, promoting gene expressions associated to adaptive stress response, neuroprotection, neuronal maintenance and longevity (60–62,56,57). In the present study, *dFOXO* expression in the PD brain was inhibited by 44% (P<0.01) during health phase and by 55% (P<0.01) during transition phase (Fig 4). However, K intervention (500 µM and 1 mM) rescued diminished *dFOXO* level significantly (P<0.05 and P<0.01) during health phase but not during transition phase. K *per se* feeding also upregulated *dFOXO* expression by 2 fold (compared to control brain) (P<0.001) during health phase but not during transition phase (Fig 4). The observation suggests that *dFOXO* expression modulation in PD and physiologic conditions is a feature of K that is restricted only to health phase. Hence, K-mediated adult life stage specific neuroprotection underlies the modulation of *dFOXO*. Further investigation revealed that in healthy transition phase brain, *dFOXO* expression is downregulated by 16% (P<0.01) as compared to that of health phase (Fig S2).

**Figure 4:**
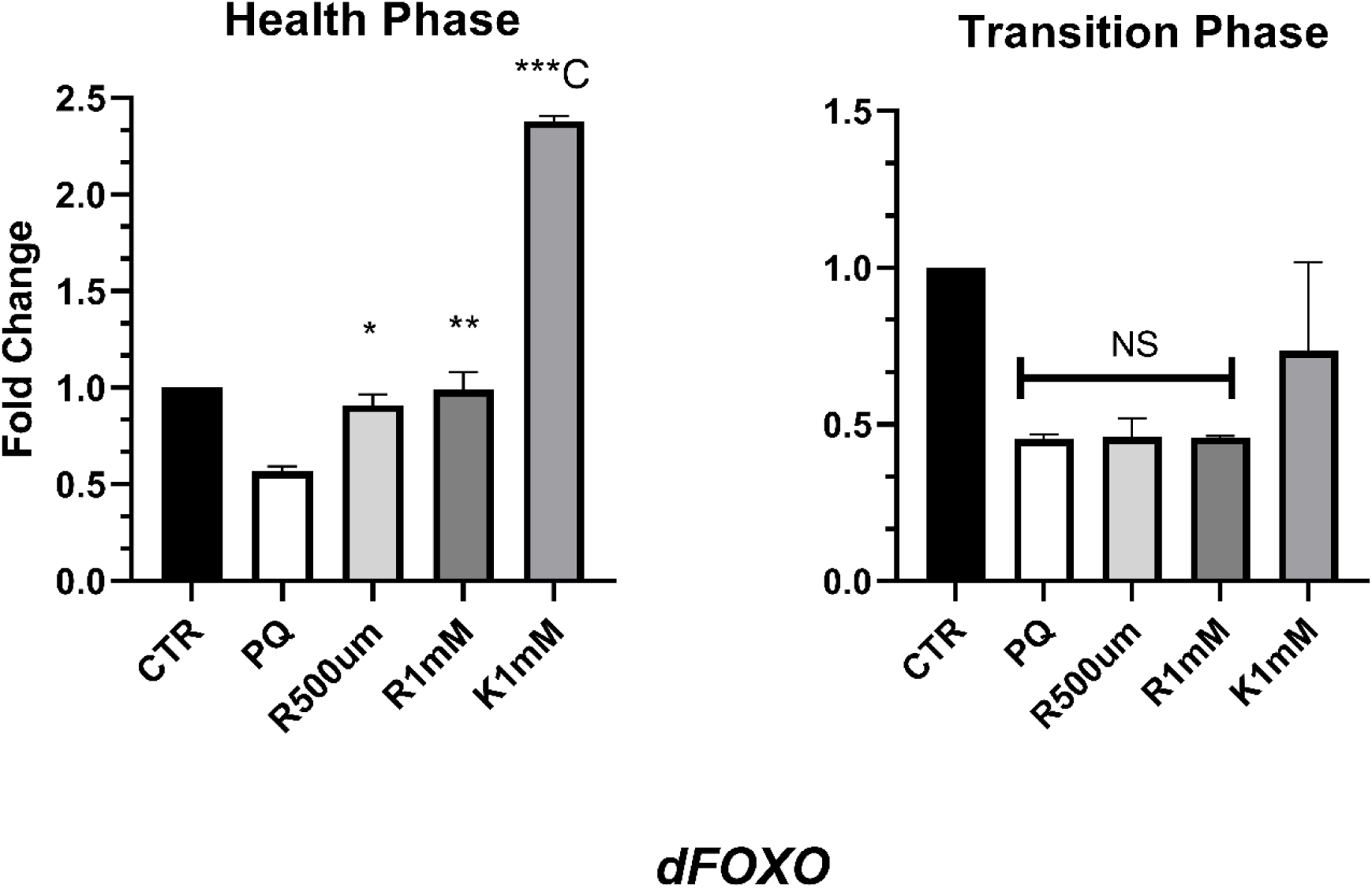
Gene Expression Analysis. K-mediated adult life stage specific differential expression of *dFOXO* involved in adaptive stress response pathway in *Drosophila* model of PD. In the PD brain *dFOXO* expression was inhibited during both health phase and transition phase. K intervention differentially regulates *dFOXO*-mediated stress response through the rescue of diminished *dFOXO* only during health phase. Significance was drawn by analysing the data of a minimum of three replicates with one-way ANOVA followed by Tukey post hoc test. [*p<0.05; **p<0.01; ***p<0.001; NS: Not significant - compared to PQ treated group], [*C p<0.05; **C p<0.01; ***C p<0.001 - compared to Control (CTR) group].

## 3. Discussion

Studies on post-mortem brains from PD patients have implicated the role of oxidative damage in the pathogenesis of PD (63–65). Evidence from multiple PD models (both in vitro and in vivo) illustrate that OS is the underlying mechanism that leads to DAergic neuronal dysfunction and demise (66). This illustrates the role of OS in DA degeneration is linked to PD.

### 3.1. Susceptibility to Oxidative Stress with Age

Protein, lipids and nucleic acids are biological macromolecules which are oxidatively altered due to ROS, resulting in genetic mutations and cellular senescence (67). In complications and pathologies associated with aging, increasing evidence for causal role of cell senescence have been experimentally shown (68). Due to stress during aging there is increase in cell senescence. This phenomenon is controlled by a number of counting mechanisms including telomere shortening, gradual increase in damage of DNA, deviation in normal oncogenes behavior, change in metabolic action and increased ROS generation (69). Such processes lead to impairment of cellular function and thus produce and accumulate ROS which in turn disturb the homeostasis, a vital factor for safeguarding the senescent phenotype. Apart from injuring DNA and damage mitochondrial activity, OS trigger the activity of p53, a gene coding for cell cycle regulator protein, which in turn induces prooxidant genes. Disturbances in redox homeostasis by generating ROS lead to deterioration of cell and impair its function and as a response to increased ROS levels, mechanisms for cellular protection are excessively activated including up-regulated expression of antioxidant enzymes (70–74) suggesting OS is intricately linked to aging and neurodegeneration.

### 3.2. Insights from Quantification of Non-Enzymatic Markers

In the present study in *Drosophila* model of sporadic PD condition, the levels of non-enzymatic OS markers such as ROS, PC, MDA, HP were increased significantly during both the phases (health and transition) of adult life. Upon K treatment sequestration of OS is evident from the significantly diminished levels of all the above markers (fig.1A-H).

While there was about 35% increase in ROS level in health stage of PQ treated flies when compared to control, in transition stage the level was increased by 5 fold suggesting stress vulnerability of the fly with age (fig.1A,B).

Increase in ROS levels and MDA an intermediate in the lipid peroxidation process upon PQ exposure has also been shown by other laboratories (75). Lipid peroxidation is a well-known example of oxidative damage in cell membranes, lipoproteins, and other lipid-containing structures. Study on postmortem PD brain have shown lipid peroxidation was increased in Parkinsonian nigra compared with other brain regions and control tissue (76). MDA levels are also found to be significantly higher in PD patients than in controls suggesting that high plasma lipid peroxidation rates might contribute as a risk factor for PD (77). PQ induced increase in the MDA level was effectively reduced with K co-treatment in both the age groups (fig.1C,D), as also shown by other reports of K ability to attenuate elevated MDA levels (78) which indicate the role of K as effective agent for neuroprotection.

There was significant protein damage in fly brain tissue as evident by the high level of PC which indicate the increased rate of protein oxidation (fig.1E,F). Protein carbonylation is an irreversible oxidative damage which leads to loss of protein function and it is considered as a widespread indicator of severe oxidative damage and disease-derived protein dysfunction. Elevated levels of PC associated with PD have been reported in postmortem PD brain tissues (79). The efficacy of K to decrease the PC levels has been reported (121,80) suggesting the neuroprotective role of K.

HP groups are powerful oxidants that can induce secondary damage by inactivating thiol-dependent enzymes essential to cell functions. Study on α-synuclein implicated in PD catalyzing the formation of hydrogen peroxide has been reported (81). Protein hydroperoxides results in modification of the structure and properties of amino acid residues. Lipid hydroperoxides are also prominent non-radical intermediates of lipid peroxidation that can often provide important mechanistic information. Increased levels of lipid hydroperoxides in Parkinsonian substantia nigra has been demonstrated (76) implicating the role of OS stress in neurodegenerative disease such as PD.

### 3.3. Estimation of Antioxidant Enzymes Activity

In *Drosophila*, SOD/CAT system is one of the intracellular protective mechanisms against ROS induced damage. Studies have demonstrated that in response to the OS generated by PQ, levels of SOD, CAT and GST are upregulated. K treatment results in the optimization of these anti-oxidant enzymes suggest lowered level of OS in the brain of K co-treated flies. Post mortem studies have shown an increased SOD activity in PD brains (82) where SOD activity was significantly associated with late PD stage. Investigation of two synthetic superoxide dismutase/catalase mimetics protecting against PQ-induced DAergic cell death in both the rat DAergic cell line and primary mesencephalic cultures in vitro and in adult mice in vivo has also been reported (83). There was significant increase in the activity of antioxidant enzymes such as SOD (fig.2A,B), Catalase (fig.2C,D) and GST (fig.2E,F). These findings substantiate well with the increased in level of ROS and other stress marker measured (fig.1A-H). K therapeutic treatment could attenuate the altered activities of these biomarkers measured in both the age groups.

The GST activity was also elevated under PD condition and K treatment markedly optimized its activity in the present study (fig.2E,F). Antioxidant role of GST in protection against oxidant toxicity and regulation of stress-mediated apoptosis has been carefully reviewed (84). GSTs are a functionally diverse family of soluble enzymes of detoxification that use reduced GSH in conjugation and reduction reactions. Depletion in GSH content and an increase in GST activity have been reported in the brains of human PD patients and also in other experimental models of PD (85,86).

Earlier investigations have also suggested the ROS to be the major factor contributing to DAergic neuronal loss in the PD brain, as a consequence of DA metabolism, low GSH, and high levels of iron and calcium in the SN*pc* (87). GSH confers the neuronal cells with multiple defenses not only to sequester but also against different types of toxic products and by products of cellular mechanism. Reduction in GSH levels as compared to control subjects in SN in PD patients have been reported (88). Depletion in GSH level may impair mitochondrial function via inactivation of complex I and increase in ROS production and the relation between decreased GSH function and severity of PD has been reported (89). In the present study K therapeutic intervention markedly rescued the GSH level in both the age groups (fig.2G,H). Efficacy of K to elevate the reduced GSH activity against Acrylamide induced toxicity has also been reported (90).

Quantification of total Thiols showed reduced activity under PD condition and K could marginally increase the total thiol levels (fig.2I,J). Thiol groups are highly reactive and susceptible to oxidation that may cause significant loss of biological activity. A critical function of cell-based thiol redox buffering systems is to protect thiol groups from oxidation and to repair those that may have become oxidized as a result of normal or aberrant cellular metabolism. Studies have shown that thiols levels are reduced under disease conditions (91).

### 3.4. Acteylcholine esterase as Neurotoxicity Marker

Neurotransmitter enzyme AChE activity in brain tissue was estimated in the present study. It was found that the enzyme activity was inhibited under PD condition in both the age groups (fig.3A,B). Such action will impair the neurotransmission process since AChE is involved in maintaining the free Acetylcholine in the synaptic region. Earlier studies have shown reduced AChE activity in several neurodegenerative diseases (92,93). AChE inhibition has been used as a biomarker for the influence of organophosphate and carbamate pesticides in invertebrate and vertebrate models. PQ exposure led to marginal inhibition of AChE activity and inhibited levels were rescued by K co-treatment in both the age groups (fig.3A,B) reaffirming the efficacy of K to attenuate the PQ induced toxicity in both the age groups. It has been proposed that exposure to low levels of pesticides lead to several neurological and neurobehavioral changes that cannot be accounted for AChE inhibition only (94,95). The particular enzyme associated with the different sub-cellular fractions can exert different effects.

The present results show that K can sequester the enhanced levels of OS not only during early phase (health stage) but importantly, also during late phase (transition stage) of adult life stage in fly PD model as seen from the above results. Further it has been observed that at higher concentration treatment, K *per se* can act as potential pro-oxidant, suggesting the necessity to properly screen the compounds before prescribing them as a therapeutic agent.

### 3.5. Oxidative Stress marker-based studies alone is not sufficient to decipher the Neuroprotective efficacy of Nutraceuticals in late onset neurodegenerative diseases such as PD

Previously our laboratory had demonstrated that neuroprotective efficacy of K is adult life stage specific (53). Many studies have demonstrated the neuroprotective efficacy of nutraceuticals using OS markers as a gold standard, considering the OS as a factor responsible for neurodegenerative diseases such as PD (96–99).

It has been demonstrated that DAergic neuroprotection of K is due to its ability to sequester oxidative stress (78,100,101). In order to understand the neuroprotective efficacy of K during health span and inefficacy during transition phase, we performed comprehensive experiments to understand K anti-oxidant efficacy with the help of multiple oxidative stress markers namely reactive oxygen species, lipid peroxide, protein carbonyl, superoxide dismutase, catalase, glutathione, glutathione S-transferase, total thiols, acetylcholine esterase activity.

We found that K can mitigate the brain OS induced by PQ exposure in health stage flies as evident from the above results. This finding confirms with several other studies of K neuroprotection employing health stage in adult *Drosophila* PD model (102). Also K could sequester the OS even in the transition stage fly brains as seen from the above study on stress markers, antioxidant and neurotransmitter enzyme activities which is an important and interesting point to note. It is shown that dietary feeding of K and other phytochemicals modulate PQ induced alteration in the expression levels of several genes that are associated with anti-oxidant and anti-aging effect in 2-3 days old flies (103). K also is reported to improve PQ induce motor deficit and replenish the decreased DA level in adult health stage (53,104). However, K is unable to rescue either the motor deficits or diminished DA level in adult transition stage (50-55 days) of *Drosophila* exposed to PQ (53). However, K could diminish the OS levels in both the fly age groups as seen from the present study, clearly suggesting that sequestration of boosted OS alone is not enough to prevent the DAergic neurodegeneration in PD. Therefore, apart from OS, there is the synergic effect of other pathways that could be responsible for the DAergic neurodegeneration in PD. Several laboratories have shown the effective role of various therapeutic compounds that modulates the altered enzymatic functions arising out of OS in model organisms (105–108), suggesting for therapeutic approach using such active compounds in NDD such as PD. However, according to the present findings it is clear that a) assessing OS levels alone b) studies in young animals i.e. belonging to health phase of adult life- is not sufficient enough to understand the neuroprotective efficacy of therapeutic molecule in late onset NDD such as PD. Studies in addition have also shown that K increases longevity when administered in the adult health stage but exerts negative result when administered in later life stages, suggesting that target molecules of K are present in the health stage (109) but may not be in transition and senescence stage. Present study reveals that OS may be a necessary factor, but alone it may not be sufficient enough for DA degeneration which is characteristic pathological feature of PD. Therefore, studies in animal models through which neuroprotective efficacy of multiple natural products being assessed based on OS markers alone may not be good enough to recommend them as neuroprotective agents.

The efficacy of K in human PD condition was tested and (80 mg/day for 9 months) on idiopathic PD patients (≥30 years of age) and the trial was unable to demonstrate its effectiveness in improving the clinical symptoms and quality of life of PD patients (110). Previously our laboratory demonstrated in a fly model of PD that DAergic neuroprotective efficacy of K is life-stage specific and this finding highlights K limitation as a therapeutic agent for late onset NDD such as PD. It is possible that the previous pre-clinical studies utilizing only adult-young animal models and/or in-vitro models that demonstrated neuroprotective efficacy of K, prompted for the need to perform clinical trial in human PD condition. This can be the possible explanation for the observation made by Kunnummakkara et al. 2023, (34) that “scientific reports based on cell culture and animal studies are often not reproducible in humans”. Hence, it is important to note that PD is a late onset disorder and therefore, implementation of ALSS model in pre-clinical studies are necessary, so as to accurately guide the course of downstream human trials for nutraceuticals and small molecules. This will help to cut short the discovery time for cutting-edge therapeutic measures against PD.

In light of the evidences, our study provides following critical insights into modeling late onset NDD such as PD and screening nutraceuticals for DAergic neuroprotection.

a. It is important and necessary to use animal models belonging to the transition stage of adult life, this is the stage of adult life during which PD sets in, for the studies relating developing a disease model and screening neuroprotective efficacy of nutraceuticals/potential therapeutic agents/drugs to late onset neurodegenerative disorders such as Alzheimers and Parkinsons disease.
b. As has been demonstrated in the present study though the K sequesters OS during both i.e health and transition phase of adult life; it fails to confer DA neuroprotection during transition phase. Hence ability to sequester the oxidative stress may be a necessity for preliminary screening of natural compounds/drugs for DA neuroprotection but not sufficient to recommend as a therapeutic agent. Hence studies need to be conducted by deciphering the genetic/molecular targets either by pathway specific fashion/genome wide molecular target analysis. Taking a cue from our study, in order to understand the neuroprotective efficacy of K we proceeded further by looking into life stage specific molecular targets through brain-specific transcriptomic and proteomic approaches, results of which will be discussed elsewhere.

### 3.6. K’s life stage-specific regulation of *dFOXO* and its role in DAergic neuroprotection

It is established in the fly model that the sole modulation of *JNK* signalling is not enough to modulate cellular physiology under stress. It was demonstrated that *JNK* signalling-mediated life span extension requires the presence of functional *dFOXO* signalling in Drosophila (57). The current study demonstrated that under the PD condition during both the adult life stages, downstream *dFOXO* expression is inhibited (Fig 4). Tas *et al*., (61) reported that inhibition of *dFOXO* signalling through *RNAi* or deletion spanning in the *dFOXO* gene (Heterozygous or homozygous condition) promotes the loss of DAergic neurons and selective loss of PAM DA neuronal cluster. It shows that *dFOXO* is necessary for neuronal maintenance which corroborates with the diminished *dFOXO* level in PD brain and neuronal dysfunction in the present study. K intervention rescued diminished *dFOXO* level only during health phase but not during transition phase (Fig 4). It was reported in a *PINK1* null fly model that overexpression of *dFOXO* and downstream genes (*SOD2*, *4e-bp*) in the thorax and brain could rescue abnormal mitochondrial phenotype, motor deficit and DAergic degeneration (56).

Further, it was also reported that K-analogue tetra-hydro curcumin mediated life span extension and stress resistance could not be imparted on *dFOXO* null flies (59). Therefore, it can be postulated from the current study that K-mediated health phase specific neuroprotection involves *dFOXO* modulation which further down the line might modulate its target gene(s) of neuroprotection.

In the present study K *per se* upregulated *dFOXO* level (compared to the control brain) only during health phase (Fig 4). This signifies enhanced *dFOXO* signalling with K under the physiologic condition. Enhancement of *JNK* signalling through *JNK* and *dFOXO* in fly models promotes enhanced longevity, better proteostatic health, delayed senescence, and resistance to neurotoxicant stress (55,57,58,59). K is reported as an early-acting inducer of longevity, as it enhances fly life span when fed at early stages or during health phase of life through dietary means (109,111). Therefore, from the current study, it can be put forward that health phase specific K-mediated modulation of *dFOXO* may also contribute to longevity under normal physiological condition during health phase.

It was also observed in the current study that with aging the *dFOXO* level is inhibited in healthy transition phase brain (Fig S2), suggesting hindrance of their regulatory system with natural aging. Further, it was reported that with senescence *dFOXO* signalling activity in the muscle and neuron declines (112). It is possible that due to such aging-associated changes in *dFOXO* regulation, K intervention fails to modulate the same in the PD brain of transition phase flies. Also, such changes may contribute to the limitation of K *per se* to modulate *dFOXO* in the physiologic condition during transition phase.

The current study clearly demonstrates that sequestration of OS by K is not good enough to promote DAergic neuroprotection. Rather it is the health phase-specific rescue of diminished *dFOXO* that may underlie ALSS neuroprotection. Resuscitation of the adaptive stress response by *dFOXO* and its possible downstream targets may contribute to DAergic neuroprotection. During transition phase such is not possible due to natural aging associated hindrances in the *dFOXO* regulatory mechanism, leading to the limitation of K-intervention during later stages of adult life. However, due to K being a multi targeting polyphenol, there are possibilities of involvement of a battery of molecular players belonging to different biological pathways in ALSS neuroprotection. Further, molecular pathways such as IIS-*mTOR/dTOR* pathway, mitochondrial dynamics, phase II antioxidant defense system, insulin signalling and metal homeostasis are also reported crucial for DAergic neuroprotection/neurodegeneration (17,113,- 116). Therefore, we are further working in that direction to uncover the possible involvement of the molecular players in multiple pathways implicated in neurodegeneration and K-mediated ALSS neuroprotection. Though figuring out the differential brain-specific regulation of certain critical pathways of neuroprotection/degeneration is a starting point, it is essential to look into differential regulation in the brain-specific transcriptome that will throw light on novel genetic players that are responsible for life state-specific DAergic neuroprotective efficacy of curcumin. This understanding at the global level will be of great support to formulate novel feeding regimen strategies, and there lies the possibility to sustain the genetic targets beyond the health phase of adult life, i.e., the transition phase.

### 3.7. Future prospects

The present study suggests that sequestration of oxidative stress is necessary but is not sufficient to promote neuroprotection during later stages of life. Other molecular pathways/targets may be involved in curcumin-mediated DAergic neuroprotection. Failure to sustain these molecular players during the later stages of life may lead to the inefficacy of curcumin (Phom, 2018). Therefore, it is necessary to look into the molecular targets of life stage specifc neuroprotection. We are looking into the life phase-specific regulation of molecular networks involved in the JNK-signalling pathway, IIS-TOR- signaling pathway, mitochondrial dynamics, phase-II anti-oxidant defense system pathway, the metal-homeostasis pathway to decipher the molecular targets of neuroprotection.

Further we are looking into the whole transcriptome to figure out novel pathways of neuroprotection.

Our study provides a critical insight into the possible reason for the important observation made through human clinical trials that despite encouraging results in animal models, therapeutic efforts to target the general state of oxidative stress failed to demonstrate an impact on the disease progression of PD. In a nutshell, employing life stage-specific animal models and, importantly, not restricting the studies only to the anti-oxidant potential of nutraceuticals is necessary and essential to carry forward the studies conducted in animal models to translate to clinical settings in humans, which is the end purpose of biomedical research.

## 4. Experimental Procedures

### 4.1. Fly Stock

*Drosophila melanogaster* of Oregon K strain was obtained from National *Drosophila* Stock Centre of University of Mysore, Mysore, Karnataka, India. They were kept in Drosophila environmental chambers (Percival, USA) at 22°C-24°C and 60% relative humidity and fed on a standard culture medium made of sucrose, yeast, agar agar and propionic acid.

Adult life-phase specific (health and transition stages) fly PD model that was developed in our laboratory (53) is employed in the present study.

### 4.2. Chemicals

Bovine Serum Albumin (A-2153), Curcumin (C-1386), Paraquat (methyl viologen dichloride hydrate, 856177), 2,7-dichlorofluorescin diacetate (D-6883), Thiobarbituric acid (T5500), Hydrogen peroxide (323381), Acetylthiocholine iodide (A5751), Quercetin (fluka, 200595), N’N’N’N-Tetramethylethylenediamine (T9281), Glutathione (G-4251), were purchased from Sigma Aldrich St. Louis, USA. Dimethyl sulfoxide (DMSO), 2,4-Dinitro phenyl hydrazine (DNPH), 1,1-dithio nitro-bi-benzoicacid (DTNB), 1-Chloro-2,4-dinitrobenzene (CDNB), Ethyldiaminetera acetic acid (EDTA), Dimethlysulfoxide (DMF), Sodiumdodecylsulphate (SDS), Ortho-pthalaldehyde (OPA), Xylenol orange, Acetone, Glacial acetic acid, Trichloroacetic acid and all other analytical grade chemicals were procured from Sisco Research Laboratory Chemicals and Merck, India. Whatman filter paper no.1 disc was used as a feeding medium in the experiment.

RNaseZAP (Ambion, Waltham, USA, Cat: AM9780), DEPC treated water (HiMedia, Thane, India, Cat: ML024), TRIzol^TM^ reagent (Invitrogen, Waltham, USA, Cat:15596026), Chloroform (Sigma-Aldrich, St. Louis, USA, Cat: c2432), Isopropyl alcohol (HiMedia, Thane, India, Cat: MB063-1l), DEPC treated water (HiMedia, Thane, India, Cat: ML024), Ethanol (MERK, Rahway, USA, Cat: 1.00983.0511), DNase (Invitrogen, Waltham, USA, Cat:18068- 015), Oligo(dT) (Invitrogen, Waltham, USA, Cat: 18418012), dNTP mix (Qiagen, Hilden, Germany Cat: 14505289), Superscript^TM^ II reverse transcriptase (Invitrogen, Waltham, USA, Cat: 18064014), PowerUp™ SYBR^®^ Green Master Mix (Applied Biosystems, Waltham, USA, Cat: A25742), 96 reactions well plate (Biorad, Hercules, USA, Cat: MLL-9601),

### 4.3. Treatment Protocol

Male flies were aged for 4-5 days (health phase) and 50-55 days (transition phase) feeding in sucrose-agar media. The flies were transferred to freshly prepared media every 3^rd^ day while aging. The flies were then transferred to vials (30mm x100mm) containing disc of Whatman filter paper no. 1 saturated with 275µl of 5% sucrose, 10mM PQ in 5% sucrose, Curcumin (500µM, 1mM) with PQ (10mM) and Curcumin in DMSO. At 24hrs of exposure, flies were frozen at -80°C. For dissection of head, an aluminium tray was positioned on the ice block and flies were placed on it. Using Carl Zeiss stereozoom (Stemi 305) microscope the flies were then dissected separating head from body with a sharp razor.

### 4.4. Total Protein Isolation from Fly Heads

100 heads were homogenized with pestle motor mixer (Argos technologies) in 175µl 0.1M Phosphate Buffer Saline (PBS) (pH 7.4). It was then centrifuged at 5000rpm for 10mins @ 4°C. The supernatant was again centrifuge under the same conditions. The resulting clear supernatant was stored at -80°C/used for biochemical assay.

### 4.5. Estimation of Protein

The Protein concentrations of the tissue homogenates were determined by the modified version of the method initially described by Bradford (117) using Bio-Rad protein assay dye reagent concentrate. The Bradford assay is a colorimetric assay for protein determination based on absorbance shift in the dye Coomassie brilliant blue-G250. Coomassie brilliant blue which is red in unbound form, on binding to protein change to stable blue form with absorbance shift from 465nm to 595nm. Since the increase of absorbance at 595nm is proportional to the amount of bound dye and thus to the amount of protein present in the sample. This can be used as a measure for the protein concentration of the unknown sample. Bovine serum albumin was used as the standard prepared in concentration range of 0.5µgP to 3.5µgP. The measurement was performed using NanoDrop 2000 (Thermo Scientific).

### 4.6. Biochemical Assays

#### 4.6.1. Reactive Oxygen Species (ROS)

Generation of ROS was determined using 2’,7’-dichlorofluorescein diacetate (DCFDA), a non polar compound that can quickly react with ROS to form the highly fluorescent compound, 2’,7’-dichlorofluorescein (DCF) after conversion to a polar derivative by intracellular esterases. DCF can be detected by fluorescence spectroscopy.50µgP of brain sample was incubated in Locke’s buffer (NaCl-154, KCl-5.6, NaHCO_3_-3.6, HEPES-5, CaCl_2_-2, glucose- 10 in mM concentrations, pH 7.4) containing 5µM DCFDA for 45 minutes at room temperature and the fluorescence was measured with excitation 480nm, emission 530nm using Cary Eclipse Fluorescence Spectrophotometer (Agilent Technologies).

#### 4.6.2. Lipid Peroxide (LP)

LP was measured with estimation of malondialdehyde (MDA) by thiobarbituric acid (TBA) reaction method. 200µgP of test samples were allowed to react in 1.5 ml of 20% acetic acid (pH 3.5), 1.5 ml of 0.8% TBA, 0.2 ml of 8% SDS (w/v) and vortex, followed by 100°C incubation in water bath for 30 minutes. The mixture was cool down to room temperature and optical density absorbance was measured at 532nm using NanoDrop 2000 (Thermo Scientific) (118).

#### 4.6.3. Protein Carbonyls (PC)

This technique involves detecting and quantifying oxidative modification of proteins. PC levels in the samples were quantified following reaction of 100µgP test samples with 2,4- dinitrophenyl hydrazine (DNPH) for 1hr. The protein is precipitated by adding 20% trichloroacetic acid, kept on ice for 5 minutes and it is then centrifuged at 3000rpm for 10 minutes at 4°C. The pellets were washed with 1ml ice cold acetone, followed by dissolving in 20mM Tris-HCl buffer (pH 7.4, containing 2% SDS, 10mM EDTA). The optical density was measured at 360 nm using NanoDrop 2000 (Thermo Scientific) and expressed as nmol carbonyl/mg protein (MEC=22,000/M/cm) (119).

#### 4.6.4. Hydroperoxides (HP)

HP levels were measured according to a previously described method using Ferrous ion oxidation xylenol orange (FOX1) reagent (100µM xylenol orange, 250µM ammonium ferrous sulphate, 100µM sorbitol, 25mM H_2_SO_4_). 100µgP of test sample was added to 1ml FOX reagent and incubated in dark at room temperature for 30 minutes. The color developed was read at 560nm in a NanoDrop 2000 (Thermo Scientific). The concentration of HP was calculated using molar extinction coefficient of 2.2 x 105 M^-1^ cm^-1^ and expressed as nmol HP/mg protein (120).

#### 4.6.5. Superoxide Dismutase (SOD)

Superoxide dismutase activity was measured indirectly by monitoring the inhibition of quercetin auto-oxidation. 100µgP of test sample was mixed with phosphate buffer (0.016 M, pH 7.8, containing TEMED- 0.8mM and EDTA-0.08 mM) followed by addition of quercetin (1.5mg/10 ml DiMethylFormamide). The rate of its auto-oxidation was monitored at 406 nm for 1 minute with 10 seconds interval using NanoDrop 2000 (Thermo Scientific). 50% inhibition of quercetin oxidation in the test sample is defined as one unit of the enzyme and activity expressed as units/mg protein (121).

#### 4.6.6. Catalase (CAT)

CAT activity was estimated by adding 25µgP of samples to phosphate buffer (0.1M, pH 7.4, containing 10 mM H_2_O_2_). H_2_O_2_ degradation was monitored at 240nm for 2 minutes with 15 seconds interval in NanoDrop 2000 (Thermo Scientific) and expressed as nmol substrate/min/mg protein (MEC=44/mM/cm) (122).

#### 4.6.7. Glutathione S-Transferase (GST)

GST activity was estimated by monitoring the conjugation of glutathione to CDNB. The reaction was initiated by addition of 40µgP test sample to phosphate buffer (0.1M, pH 6.5) containing 0.5mM EDTA, 0.075mM CDNB, 0.05mM GSH. The increase in optical density was observed for 5 minutes at 340nm using NanoDrop 2000 (Thermo Scientific). The activity was expressed as nmol conjugate formed/min/mg protein using MEC=9.6 mM^-1^ cm^-1^ (123).

#### 4.6.8. Reduced Glutathione (GSH)

GSH activity was determined based on fluorimetric method using Ortho-pthalaldehyde (OPA). 50µgP of sample protein was mixed with 5% tricarboxylic acid (TCA) and centrifuge at 3000rpm for 5 minutes at 4°C. The supernatant was added to a reaction mixture of buffered formaldehyde (1:4 (v/v) 37% formalin: 0.1M Na_2_HPO_4_). 900µl of Sodium phosphate buffer (0.1M, 5mM EDTA, pH 8.0) was added to each tube followed by 100µl OPA. Then after dark incubation for 30 minutes at room temperature, the fluorescence was measured at excitation and emission wavelengths of 345 and 425 nm respectively using Fluorimeter (Agile Life technologies) (124).

#### 4.6.9. Total Thiols

The procedure is based on the reaction of the thiol with 5,5’-dithiobis 2-nitrobenzoic acid (DTNB) to give the mixed disulfide and 2-nitro-5-thiobenzoic acid (TNB) which is quantified by the absorbance of the anion (TNB^2-^) at 412 nm. Total thiols activity was determined by taking 25µgP of test sample containing Tris buffer (pH 8.0) reacted by adding 25µl DTNB and 1775µl methanol. The reaction was incubated in dark for 30minutes with occasional mixing, followed by centrifugation at 3000rpm for 5 minutes at 4°C. The supernatant was used to measure the ability of thiols to oxidize DTNB using spectrophotometer at 412 nm in NanoDrop 2000 (Thermo Scientific). Total thiols activity was then expressed as nmol/mg protein (125).

#### 4.6.10. Acetylcholinesterase (AChE)

AChE efficiently catalyze the hydrolysis of acetyl-thiocholine sulfur analogs of its natural substrate, acetylcholine. AChEactivity was determined by taking 25µgP test sample containing in phosphate buffer (0.1M, pH 8.0) and DTNB (10mM). To the mixture, 20µl of acetylthiocholine iodide (ATCI, 78 mM) was added. The reaction absorbance was monitored at 412 nm for 5 minutes with 30 seconds interval in a NanoDrop 2000 (Thermo Scientific). AchE activity was then expressed as n mol substrate hydrolyzed/min/mg protein (126).

### 4.7. Gene Expression Analysis Using the Quantitative Real Time PCR (qRT PCR)

The methodologies pertaining to gene expression analysis utilizing qRT-PCR is partially described in Das *et al*, 2021 (127).

#### 4.7.1. Total mRNA Isolation from Fly Heads

In brief, post-exposure each group of flies were frozen and heads were decapitated. 50 heads per fly group were used to extract total RNA extraction using Guanidium thiocyanate phenol chloroform extraction or the TRIzol method. Tissues were homogenized with 1 mL of TRIzol and centrifuged (At 12g for 15 mins at 2°C) to discard the tissue debris. Supernatant was collected and incubated at room temperature for 5 mins. Phase separation by chloroform (200 µL was added and vigorously shaken followed by incubation at room temperature for 2-3 mins) was opted to be the best way to isolate aqueous solution containing RNA from the homogenate. The tubes are centrifuged at 12g for 15 minutes at 2°C. Colourless upper aqueous phase was collected and RNA was eluted and precipitated out from the aqueous solution by adding chilled Isopropyl alcohol (500 µL was added and incubated at room temperature for 10 minutes followed by centrifugation at 13g for 15 min at 2°C). The pellet of RNA was washed with 1 mL of 75% ethanol to remove salt content from TRIzol reagent. Pellet was air dried and resuspended in 30-35 µL DEPC treated water (Incubated for 10 minutes at 55° to 60 °C to dissolve the pellet). RNA was quantified and stored in -80°C freezer till further use. Precaution taken to avoid RNase contamination, so every instrument and working space was cleaned with RNase ZAP prior to the experiment.

#### 4.7.2. First Strand cDNA Synthesis

RNA was quantified with pedestal method using Nanodrop (Thermo-Fisher scientific). 1 ug of RNA from each group is to be synthesized into single stranded cDNA. The required volume of RNA which contains 1 ug of the biomolecule was taken and treated with DNase with DNase buffer (1 µL each) to avoid DNA contamination. Volume makeup for each sample was done by adding DEPC water to obtain a uniform volume of 10 µL. Sample was incubated 15 mins at room temperature. After which 1 µL of EDTA was added followed by heating at 65°C for 10 minutes to deactivate the DNase. Tubes were puff centrifuged and Oligo DT and dNTP mix (1 µL each) were added followed by heating at 65°C for 5 minutes to promote annealing. Tubes were chilled immediately and then puff centrifuged. First strand buffer (4 µL) and 0.1 M DTT (2 µL) were added followed by heating at 42°C for 2 minutes to prepare the RNA template for first strand cDNA synthesis. 1 µL of Superscript II reverse transcriptase was added and incubated at 42°C for 50 minutes. The reaction was deactivated by incubating the tubes at 70°C for 15 minutes. Tubes were puff centrifuged again and now the first strand cDNA is ready which is stored in -20°C freezer till further use.

#### 4.7.3. Gene Expression Analysis

The expression analysis with Real time PCR principles reaction was carried out in Applied Biosystem (ABI) Step-One Plus thermal cycler (Thermo fisher scientific), using Power Up SYBR green master mix (containing STBR green as reporter dye). cDNA was diluted with DEPC water with a factor 5 for the reaction. Reaction volume was considered 10 µl which contains 1 µl of diluted cDNA, 8.6 µl of SYBR green and 0.2 µl of forward and reverse primer specific to the internal control *RP49* and target gene *dFOXO*. Reaction plate was loaded accordingly as described in Das *et al*, 2021 (127). The thermal cycler protocol for the qRT-PCR is as follows Amplification (Amp.) protocol: Holding stage @ 95°C for 10 mins Amp. Cycle: (95°C for 15 secs followed by 60°C for 1 min) X 40 Melting Curve analysis: Quickly ramped up to 94°C followed by cooling at 60°C (Already pre set in user interface software and can be directly selected)

Following primers were specific to the molecular targets were used

*RP49* (CG7939): F- AGGGTATCGACAACAGAGTG, R- CACCAGGAACTTCTTGAATC

*dFOXO* (CG3143): F- TCGAGTGCAATGTCGAGGAG, R- AGCGGTATATTGATGTCCAGCAG

The C_T_ values were obtained and the relative fold change to the Control sample was measured by the 2^-ΔΔCt^ method as described by Livak and Schmittgen (2001) (128).

### 4.8. Western blotting

#### 4.8.1. Protein Extraction from *Drosophila* brains

From each of the treatments 50 fly heads were homogenized in 160ul of RIPA buffer (50mM Tris HCL, 1% Triton, 0.5% sodium deoxycholate, 150mM NaCl, 0.1% SDS, 2mM EDTA) with protease inhibitor cocktail. Homogenates were then sonicated for 20sec (with pulse of 10sec and amplitude at 30%) using Qsonica sonicators (from OHIO industries). The samples were centrifuged at 13,000rpm for 5min at 4⁰C. Supernatant was re- centrifuged at 13,000rpm for 5min at 4⁰C. Lysates were stored at -80⁰C until quantification was performed.

#### 4.8.2. Protein quantification

protein quantification was performed using the Bio-rad RCDC assay reagents (cat no. 500- 0120). Bovine serum albumin at concentration of 1mg/ml (cat no. A-2153, Sigma) was used as the standard and 5ul of the extracted protein lysates were used for quantification of the samples. Absorbance was read at 750nm wavelength.

#### 4.8.3. SDS gel casting

SDS-polyacrylamide gels were cast using the TGX stain-free fast cast acrylamide kit, 10% from Bio-Rad (cat no 161-0183TA) of 1.5mm thickness.

#### 4.8.4. Bio-rad proprietary method for western blotting

1. Prepare resolving gel solution by mixing equal volumes of resolver A and resolver B solution (as described by the manufacturer).
2. Use casting stand to stabilize the glass plates during casting.
3. Add required volume of TEMED and freshly prepared 10% APS to the combined resolver and mix well. Steadily dispense the solution into the glass plates. Fill the cassette to 1cm below the bottom of the teeth of comb.
4. Prepare stacking gel acrylamide solution by combining equal volumes of stacker A and stacker B solution (as described by the manufacturer).
5. Add the required volume of TEMED and 10% APS to the stacker solution and mix well. Pipet the solution in the middle of the cassette, filling to the top of short plate. Apply slowly and steadily to prevent mixing with resolving solution. Align and insert the comb in the cassette. Allow the gel to polymerize for 30-45 min before electrophoresis.
6. 40ug of each sample lysate were mixed with 20ul of sample buffer (0.5M Tris HCl pH6.8, 10%SDS, glycerol, 0.1% bromophenol blue, β mercaptoethanol) in a total volume of 40ul and denatured for 5min at 95⁰C.
7. Bio-Rad prestained plus protein dual color standard (161–0374) was used as the protein ladder marker.
8. A 10X stock of running buffer (Tris, Glycine, and SDS) was used for preparing 1X running buffer.
9. The gel was run at a current of 20mAmp at room temperature using Bio-Rad powerpac basic power system.
10. Before electro blotting, the stain free gels were scanned and activated (2.5min) using the Bio-Rad fluorescent documentation system.
11. PVDF membrane (Bio-Rad 162-0174) was used after activation by methanol (Merck) before setting up the transfer sandwich.
12. Transfer was carried out at a voltage of 90V for the duration of 90min using chilled 1X transfer buffer (Glycine, Tris and methanol) with continuous stirring using magnetic bead. To create a cold temperature condition the transfer tank placed inside a bucket filled with ice.

#### 4.8.5. Membrane blocking and Antibody treatment

Post transfer the PVDF membrane was scanned using the Bio-Rad fluorescent documentation and incubated in the blocking buffer of 5% BSA in 1X TBS-T (0.05%) for the duration of 90min at room temperature with gentle rocking. Rabbit polyclonal to Catalase (ab16731), SOD 2 (ab13534), SOD 1 (ab13498) were used in dilutions of 1:2500, 1:2500 and 1:7500 respectively. The membranes were incubated at 4°C for 24hrs. Post primary antibody incubation, the membranes were washed 3X in 1X TBS-T (0.05%) for 15min at room temperature. The secondary antibody Goat anti rabbit HRP (ab 205718) was used in a dilution of 1:5000 and the membranes was incubated for 2hr at RT. The membranes were washed 5X in 1XTBS-T (0.05%) for 15 min and developed using Clarity western ECL substrate (Bio-Rad 170-5060). Scanning was performed with Bio-Rad fluorescent documentation system. For internal control, Actin (ab8224) mouse monoclonal antibody in the dilution of 1:4000 was used with incubation at 4⁰C for 24hrs and secondary Goat anti mouse HRP (abcam 205719) was used in a dilution of 1:5000 with 2hrs incubation at RT. PVDF membrane was stripped and re-probed for Actin internal control. The membrane was washed in stripping buffer @ RT for 10mins vigorously followed by washing with 1X PBS @ RT for 15mins and final wash with 1X TBS-T (0.1%) @ RT for 10mins. The membrane was then incubated with blocking buffer proceeded by antibody incubation as stated above.

#### 4.8.6. Data analysis

Data analysis was performed using ImageLab 5.2.1 version software.

#### 4.8.7. Methodology

Data analysis was done using the ImageLab 5.2.1 version software.

Normalization with internal control Actin (Fig. S3)

1. Open the image of target protein blot and internal control blot
2. Select lanes by manually selecting the number of lanes. Adjust and resize the lanes to the band of interest on both the target protein blot and internal control blot.
3. Click on bands and select detect bands for both the blots.
4. Bands are highlighted in the respective lanes.
5. Select each blot, go to file, select export and select lane and band file to excel. This will give the volume of each bands in their respective blots.
6. In a new excel sheet, calculate the expression of the band for each of the lane in the target protein blot as per formula: **expression= target band volume/ internal control volume.** Calculate the normalized expression of target protein by dividing with control expression into all the groups.

## Supporting information

This article contains supporting information. (Figure S1, S2,S3)

## Acknowledgement

This research was supported by the Department of Biotechnology (DBT), India (R&D grant no. BT/405/NE/U-Excel/2013) awarded to SCY. Part of this work was presented by LP at Sakura science exchange program (organized by Japan Science and Technology Agency) held at Tsukuba Science City, Japan during October 14-21, 2018; International Conference on Nutraceuticals and Chronic Diseases (INCD-2019), 23-25 September, 2019, held at the Indian Institute of Technology (IIT-G), Guwahati, India by PM, AD; *108* Indian Science Congress. (invited talk) Nagpur, India. 3-7 January, 2023 by SCY. LP, PM, AD received JRF and SRF from DBT (Department of Biotechnology), India.

## Author contribution

LP: performed assays relating to oxidative stress markers, AD: performed experiments relating to transcript expression, PM: performed experiments relating to protein expression and western blotting. LP, PM, AD drafted the respective sections of the ms. SCY:

Conceptualization, methodology, funding acquisition, review and editing of the ms, and supervision.

## Abbreviations used

(AChE): Acetylcholinesterase
(ALSS): Adult life stage-specific
(CAT): Catalase
(K): Curcumin
(R_500µM /1mM_): Curcumin_500µM or 1mM_ cotreated with Paraquat
(K_500µM /1mM_): Curcumin_500µM or 1mM_ *per se*
(DA): Dopamine
(DAergic): Dopaminergic
(GST): Glutathione S-Transferase
(HP): Hydroperoxides
(LP): Lipid Peroxide
(MDA): Malondialdehyde
(NDD): Neurodegenerative disorders
(OS): Oxidative stress
(PQ): Paraquat
(PD): Parkinson’s Disease
(PC): Protein Carbonyls
(ROS): Reactive Oxygen Species
(GSH): Reduced Glutathione
(SOD): Superoxide Dismutase

