## Supplementary material for "Sequestration of oxidative is necessary but not sufficient enough to conclude dopaminergic neuroprotective efficacy of curcumin: Insights from ALSS *Drosophila* Parkinson’s disease model": Supplemetal Fig

### Title

# S-1

#### A. Catalase

**HEALTH PHASE**

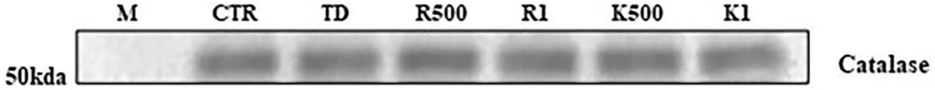


| Mol. Wt | CTR | TD | R500 | R1 | K500 | K1 |
|---|---|---|---|---|---|---|
| 55.7 kda | 1 | 0.80648 | 0.90488 | 0.87773 | 0.82438 | 0.736447 |

**TRANSITION PHASE**

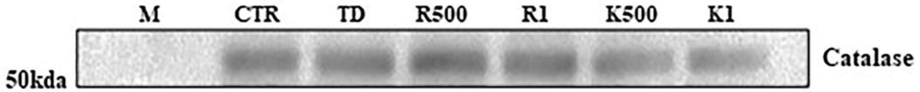


| Mol.Wt | CTR | TD | R500 | R1 | K500 | K1 |
|---|---|---|---|---|---|---|
| 55.7 kda | 1 | 0.91305 | 0.98557 | 0.96735 | 0.73732 | 0.514075 |

#### B. SOD 1

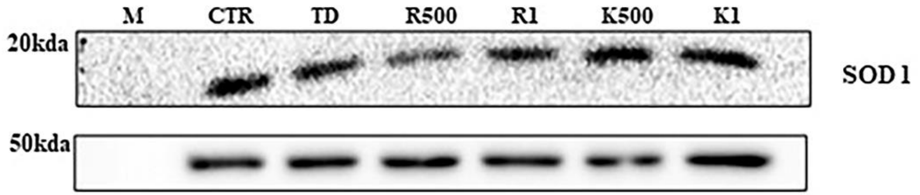


| Mol. Wt | CTR | TD | R500 | R1 | K500 | K1 |
|---|---|---|---|---|---|---|
| 18 kda | 1 | 0.71257 | 0.60129 | 0.68794 | 1.096531 | 0.72340 |

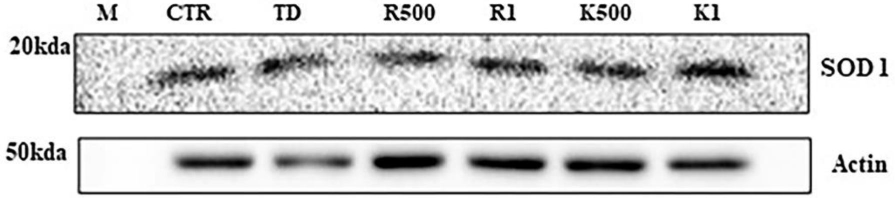


| Mol. Wt | CTR | TD | R500 | R1 | K500 | K1 |
|---|---|---|---|---|---|---|
| 18 kda | 1 | 1.07756 | 0.68349 | 0.81220 | 0.728557 | 1.17637 |

#### C. SOD 2

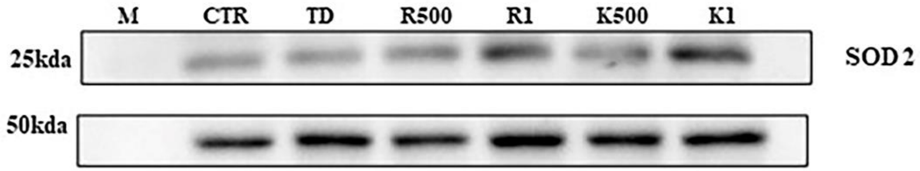


| Mol. Wt | CTR | TD | R500 | R1 | K500 | K1 |
|---|---|---|---|---|---|---|
| 25 kda | 1 | 0.86925 | 1.08379 | 1.37314 | 0.87090 | 1.65293 |

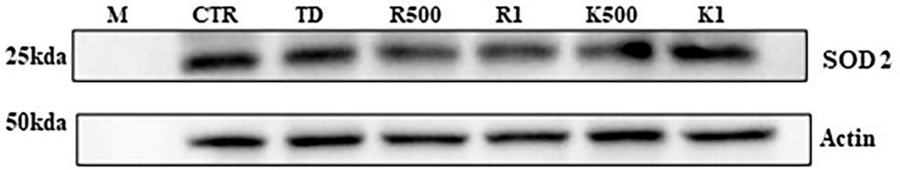


| Mol. Wt | CTR | TD | R500 | R1 | K500 | K1 |
|---|---|---|---|---|---|---|
| 25 kda | 1 | 0.84599 | 0.82089 | 0.76219 | 0.74039 | 0.86283 |

**S1:** Characterization of protein expression levels of brain Catalase (A), SOD1 (B) and SOD2 (C) through SDS page followed by western blotting:  Analysis illustrates that in PD brain A) catalase levels are reduced (Health phase: 10% and Transition phase: 20% and K cotreatment confers partial rescue only in Health phase); B) SOD 1 (Health phase: 30% reduction and Transition phase: no change, and K cotreatment did not rescue) C) SOD2 expression reduced (Health phase: 10% and Transition phase: 10% but K rescues in Health phase only).

# S-2

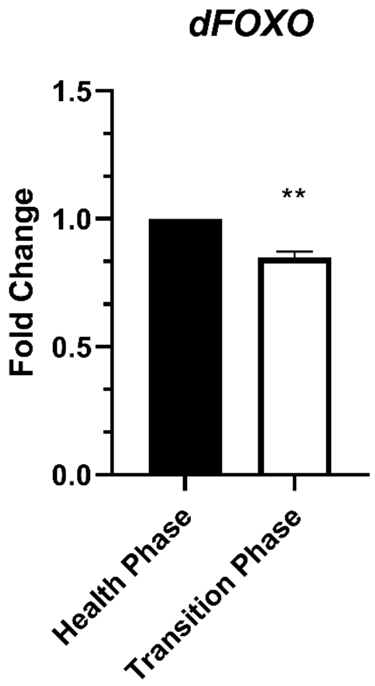


**S2:** Brain-specific life phase associated changes in expression of *dFOXO*. Natural aging in brain inhibits *dFOXO*-mediated adaptive stress response, as expression level of *dFOXO* was diminished in transition phase brain as compared to that of health phase.

# S-3

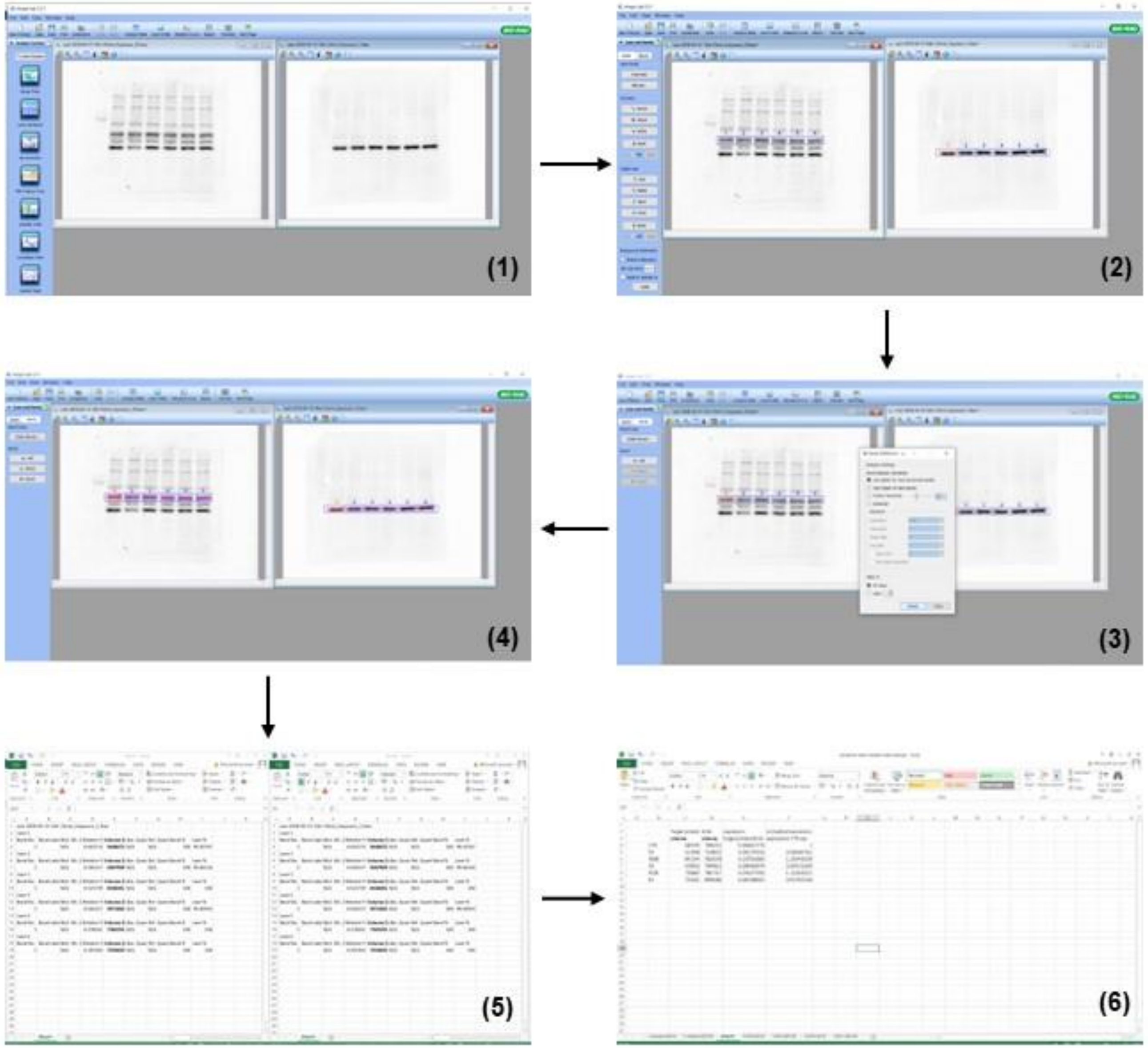


**S3:** Showing the analytical methodology behind the quantification of translate expression using BIORAD Chemidoc (whole protein normalization method)
